# Probing the polarity of spontaneous perisomatic GABAergic synaptic transmission during epileptogenesis in the mouse CA3 circuit in vivo

**DOI:** 10.1101/630608

**Authors:** Olivier Dubanet, Arnaldo Ferreira Gomes Da Silva, Andreas Frick, Hajime Hirase, Anna Beyeler, Xavier Leinekugel

**Affiliations:** University of Bordeaux, INSERM U1215, Neurocentre Magendie, 33077 Bordeaux, France; Center for Translational Neuromedicine, University of Copenhagen

**Keywords:** GABAergic transmission, inhibition, epilepsy, hippocampus, interneurons

## Abstract

Several studies suggest a contribution of reversed, excitatory GABA to epileptogenesis. But GABAergic transmission critically depends on the very dynamic combination of membrane potential, conductance and occurrence of other synaptic inputs. Taking this complexity into account implies measuring the postsynaptic responses to spontaneously occurring GABAergic events, in vivo, without interfering with neuronal [Cl^-^]i. Because of technical difficulties, this has not been achieved yet. We have overcome this challenge by combining in vivo extracellular detection of both optogenetically-evoked and spontaneously occurring unitary inhibitory postsynaptic field-potentials (fIPSPs), with the silicon probe recording of neuronal firing activity, with single cell resolution. We report that isolated acute seizures induced a global reversal of the polarity of CA3 hippocampal GABAergic transmission, shifting from inhibitory to excitatory for a duration of several tens of seconds before returning to normal polarity. Nevertheless we observed this reversed polarity only in the post-ictal period during which neurons (including GABAergic interneurons) were silent. Perisomatic inhibition was also affected during the course of epileptogenesis in the Kainate model of chronic epilepsy. One week after Kainate injection, the majority of pyramidal cells escaped inhibitory control by perisomatic GABAergic events. Besides, we did not observe a reversed polarity of fIPSPs, but fIPSPs provided time-locked excitation to a minor subset of CA3 pyramidal neurons. Beside methodological interests, our results suggest that subtle alterations in the regulation of [Cl^-^]i and perisomatic GABAergic transmission already operate in the hippocampal circuit during the latent period that precedes the establishment of chronic epilepsy.

## Introduction

GABAergic inhibition relies on ionotropic GABA_A_ and metabotropic GABA_B_ receptors. The fast and time-locked inhibition considered essential for neuronal coding is the one mediated by GABA_A_ receptors (GABA_A_ R), permeable to chloride ions (Cl^-^). In normal conditions, neurons maintain a low intracellular concentration of chloride ([Cl^-^]i), so that the reversal potential of Cl^-^ mediated currents is more negative than the resting potential (1). GABA release therefore generates an influx of Cl^-^, with an hyperpolarizing effect on membrane potential, and inhibition of neuronal discharge.

Recent publications have proposed the implication of defective inhibition in the etiology of various pathologies including epilepsy, depression, schizophrenia, Down Syndrome or Autism Spectrum Disorders (2–9). In vitro and in vivo experiments have suggested that defective function of the K-Cl cotransporter KCC2, that would normally extrude Cl^-^ from the cell, was responsible for a reversed polarity of GABAergic transmission from inhibitory to excitatory, participating in epileptogenesis in animal models (10–21) as well as in human tissue resected from epileptic patients (22–24). But one major difficulty in the study of GABAergic inhibition is to avoid artefactual interference with the native Cl^-^gradient, a main determinant of GABA_A_ R mediated inhibitory function. Perforated-patch clamp recordings with the compound antibiotic gramicidin provide electrical access to the cell through the formation of pores in the membrane, permeable to cations but impermeant to Cl^-^. Perforated-patch recordings with gramicidin have been successfully used in vitro to record GABAergic synaptic currents, preserving intracellular Cl^-^ (10, 11, 16, 21, 25). But Cl^-^ gradient has also been shown to be affected by the slicing procedure (26–28), questioning the claims for excitatory GABAergic transmission based on in vitro recordings (27).

In vivo treatment with the diuretic bumetanide, which also reduces the accumulation of Cl^-^ ions in neurons through the inhibition of the Cl^-^ intruder NKCC1, was found to reduce behavioural abnormalities in rat and mouse models of Autism Spectrum Disorders(5) and seizures in animal models of epilepsy (19, 29–33). But bumetanide has a limited bio-availability in the brain due to poor blood brain barrier penetration (34–36), and it is finally unclear if its action in vivo was indeed due to the restoration of neuronal Cl^-^ gradient and inhibitory GABAergic transmission.

Direct investigation of the polarity of GABAergic transmission in vivo puts serious constraints in terms of experimental access, and the evaluation of responses to spontaneous synaptic GABA_A_ R activation is challenging. The ideal method would fully preserve the exact spatio-temporal interactions between membrane potential, conductance and ongoing synaptic inputs (both excitatory and inhibitory), which natural dynamics are highly complex and yet critical for the net inhibitory or excitatory effect of GABAergic transmission (18, 25, 37–40). Adapting to in vivo conditions the extracellular recording of unitary GABAergic postsynaptic potentials (field IPSPs) previously described in vitro (41–43), we could evaluate the polarity of perisomatic GABAergic signaling during spontaneous activity in the intact brain. Our results provide direct evidence that perisomatic GABAergic transmission exert depolarizing actions and time-locked excitation of CA3 pyramidal neurons in acute and chronic mouse models of epilepsy, and that the majority of pyramidal cells escape inhibitory control by perisomatic GABAergic events during epileptogenesis. We believe that this demonstration will pave the way for the investigation of Cl^-^ homeostasis and inhibitory function in vivo, in physiological conditions and in situations in which alteration or even reversal of GABAergic transmission is hypothesized to occur. It also constitutes an invaluable tool to quantify the actual in vivo efficacy of drugs designed to modulate [Cl^-^]i and restore physiological GABAergic inhibition, thereby meeting high clinical and therapeutic expectations.

## Results

### Extracellular recording of perisomatic field-IPSPs and time-locked inhibition of CA3 pyramidal cell firing in vivo

Non-invasive recording of the dynamics of GABAergic synaptic transmission in vivo is a challenge. Previous work in vitro has suggested that perisomatic inhibitory postsynaptic signals originating from individual interneurons (basket and chandelier cells) could be recorded as extracellular field potentials (fIPSPs) from their target pyramidal-cell population (41–43). This approach offers several main advantages. First, extracellular recordings preserve native [Cl^-^]i and other cytoplasmic factors critical for GABAergic transmission (44). Second, the polarity of the field events indicates the direction of the current elicited in the target population of pyramidal cells, thereby addressing the question of depolarizing vs hyperpolarizing polarity of GABAergic postsynaptic signaling. And third, combined with units recording and spike sorting of neuronal discharges, extracellular detection of population fIPSPs provides a time reference to investigate the net effect of GABAergic perisomatic synaptic signals on the firing of the target neurons with single cell resolution, allowing a direct quantification of the efficacy of inhibition or excitation provided by perisomatic GABAergic inputs.

In order to evaluate the efficiency and polarity of perisomatic GABAergic transmission in the hippocampus in vivo, we have therefore inserted silicon probes into the dorsal CA3 pyramidal layer of urethane-anesthetized mice, recording spontaneous local field potentials and unitary neuronal activity. As illustrated in Fig. 1, spontaneous activity from the CA3 pyramidal layer included action potentials (units) and positive local field events with a profile similar to that of fIPSPs previously described in vitro as the GABA_A_ receptor mediated postsynaptic potentials originating from PV interneurons(43). Accordingly, these events were characterized by a fast rise time and a slow decay (time to peak, 2.01±0.53ms, exponential decay time-constant, 3.65±0.05ms, n=8 mice), and showed the pharmacological profile of GABA_A_ R-mediated IPSPs. As illustrated in Fig. S1b, fIPSPs persisted (even though with reduced frequency) in presence of locally injected glutamatergic antagonists (frequency of occurrence, 16.3±5.6% of control in APV+DNQX, 5 and 2mM respectively, 200 to 300nl), at a dose that efficiently abolished glutamatergic transmission as indicated by the loss of the LFP response evoked by the contralateral electrical stimulation of CA3 inputs. In contrast, fIPSPs were fully blocked by the GABA_A_ R antagonist bicuculine (0% of control in presence of bicu 100μM, 200nl, with or without APV+DNQX, n=3 mice, 1 way ANOVA, F(3)=872.37, p<0.001). The decay kinetics of fIPSPs were prolonged by the GABA_A_ modulators diazepam (2mg/kg, IP) and zolpidem (10mg/kg, IP). Because fIPSPs amplitude is directly affected by resting membrane potential, an uncontroled parameter in our extracellular recording conditions, we have restricted our analysis to the decay kinetics of fIPSPs. As illustrated in Fig. 1c, zolpidem steadily increased spontaneous fIPSPs decay time-constant by about 180% (from 4.66±0.36ms to 13.07±4.18ms, n=5 mice, p<0.01, two-tailed paired t-test, t(4) =-4.62), in contrast to the very modest effects of diazepam (around 15% increase, from 3.65±0.05ms to 4.19±0.06ms, n=3 mice, p<0.01, two-tailed paired t-test, t(2)=9.13). This pharmacological profile actually suggests that among PV interneurons, fIPSPs originate preferentially from basket cells rather than from chandelier cells or bistratified neurons (45, 46). Also as observed in vitro and expected from perisomatically projecting interneurons (basket and chandelier cells), in vivo fIPSPs provided very powerful time-locked inhibition of local pyramidal cells. Accordingly, multi-unit pyramidal cell activity was almost fully interrupted by fIPSPs, as observed in the traces shown in Fig. 1b and in the trough that follows fIPSPs in the peri-event time histogram (PETH) shown in Fig. 1d-e. Beside the peak firing that precedes fIPSPs and probably reflects the recruitment of perisomatic inhibition by pyramidal cells discharge(43), most of the 389 putative pyramidal cells obtained from 18 mice were significantly inhibited by fIPSPs (71.1±12.9% of the neurons tested, n=18 mice), resulting in a population firing reduced by 83.5±10.4%, returning to baseline values after 10.2±3.5ms (n=18 mice).

**Figure 1:**
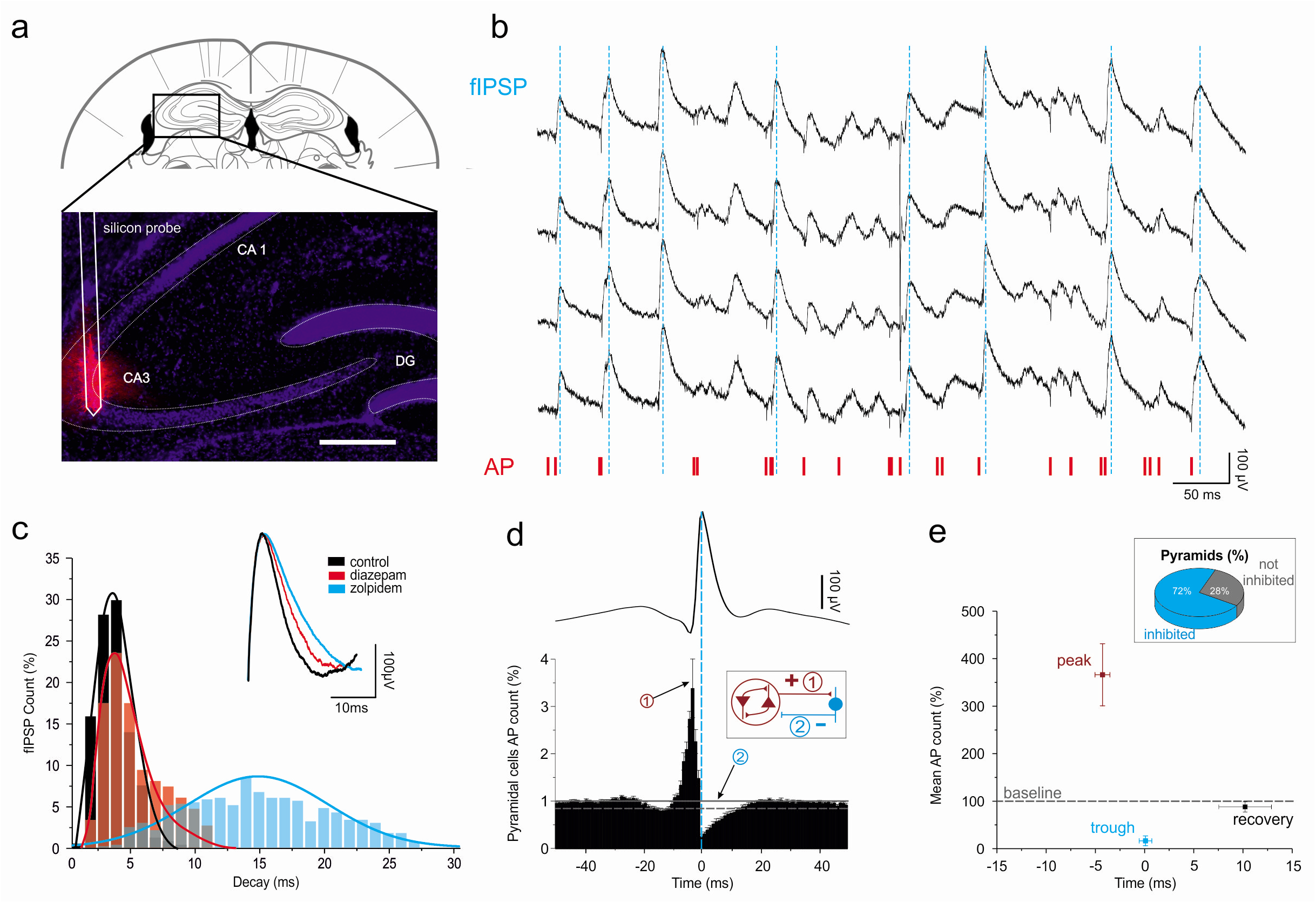
Spontaneous extracellular field inhibitory post-synaptic potentials (fIPSPs) in vivo. **(a)** Schematic of electrode track and histological verification of recording location within the CA3 hippocampal region (Dil-labeled silicon probe in red, DAP1 staining in blue). **(b)** Example trace of spontaneous wide-band (0.1 Hz-9KHz) field-recording activity in the CA3 pyramidal layer, displaying 4 channels (10-20μm distance between recording sites) of the same shank of a 2-shanks Neuronexus Buzsakil6 silicon probe. Field-events of positive polarity (upward deflections), fIPSPs (peaks, **dashed blue lines**). Fast downward deflections, action potentials (AP), also labeled below as raster display (**red ticks**, multi-unit activity). Note the transient interruption of neuronal firing after fIPSPs. **(c)** Comparison of ñPSP decay (top, superimposed average traces normalized on amplitude, below, histogram distributions of fIPSPs decay times) in control (**black**), in presence of zolpidem (**blue**) or diazepam (**red**). Note prolonged flPSP decay in presence of zolpidem (mean± SD, n=5 mice) and diazepam (mean± SD, n=3 mice); ** p<0.01. **(d)** Peri-event time histogram (mean and SD, n=18 WT mice, time bin 1ms) between all the spikes discharged by putative pyramidal cells and fIPSPs (reference, peak-time), shown as average trace on top at the same time scale and aligned on peak (**dotted blue line**). Baseline mean and 2SD are respectively shown as **plain and dotted horizontal grey lines**. Note increased pyramidal cell discharge in the 10 ms prior to ŨPSP, and the strong inhibition (trough, max inhibition) that follows. **Inset**, schematic illustration of the recurrent excitatory/inhibitory loop: (**1**) the collective discharge of interconnected pyramidal cells activates the discharge of local interneurons, resulting (**2**) in perisomatic flPSP and inhibition of pyramidal population activity. **(e)** Diagram illustrating firing rates (mean ± SEM, same 18 mice as in d; baseline level = 100%, calculated from −50 to −30ms) at **peak** (bin of maximum firing in the 10ms prior to fIPSPs), **trough** (bin of minimum firing in the 4ms post-fIPSPs), and **recovery** (last of post-fIPSPs time bin below baseline - 2SD) time points. **Inset**, proportion of pyramidal cells (n=18 mice, for a total of 389 cells) with a significant post-fIPSPs trough (ie inhibited vs not-inhibited neurons).

### Perisomatic fIPSPs specifically originate from PV interneurons and provide homeostatic redistribution of individual pyramidal cells firing rates in the CA3 hippocampal network

In order to verify that fIPSPs actually originate from PV interneurons, we recorded the extracellular field-potential responses to the optogenetic activation of PV interneurons. PV-Cre mice were injected with the DIO hChR2 (E123T/T159C)-EYFP construct in the CA3 region in order to express the cationic actuator ChR2 specifically in PV interneurons. PV immuno-staining of the infected mice showed excellent specificity, with 96 to 100% (98.3±2%, n=3 mice) of the infected (GFP-positive) cells being also PV-positive (Fig. S2a). As illustrated in Fig. 2, brief (2ms) local optogenetic activation of PV interneurons did evoke reliable fIPSPs, together with powerfull time-locked inhibition of local pyramidal cells (n=33 pyramidal cells from 5 mice). Compared to spontaneous fIPSPs, evoked fIPSPs (n=5 mice) had a longer decay time constant (5.45±1.15ms for evoked vs 4.11±0.73ms for spontaneous, t(4)=2.93, p<0.05), faster rise time (1.54±0.15 vs 1.96±0.15ms, t(4)=-3.38, p<0.01) and larger amplitude (287.5±40.53μV vs 218.1±15.48μV, t(4)=-3.19, p<0.05). As illustrated in Fig. S1, the success rate of PV-interneurons optogenetic stimulation to evoke fIPSPs was not significantly affected by the local presence of the glutamatergic receptor antagonists APV and DNQX, but evoked fIPSPs were fully (and reversibly) blocked by the GABA_A_ R antagonist bicuculine (success rate 81.5±7.1% in control, 91±11.6% in APV+DNQX, 0% in bicu, with or without APV+DNQX, 1-Way ANOVA, F(3) = 162.15). PV interneuron firing is therefore sufficient to evoke fIPSPs. In order to know whether other types of interneurons also significantly contributed to the generation of perisomatic fIPSPs during spontaneous CA3 activity, PV-Cre mice were injected with the FLEX-rev::PSAML141F,Y115F:GlyR-IRES-GFP construct to express the inhibitory pharmacogenetic actuator PSAM-GlyR specifically in PV interneurons. PV immuno-staining of the infected mice showed that in addition to high specifity (99±1 %, n=3 mice), 98±1 % (n=3 mice) of the PV-positive cells in the region of recording were also GFP-positive (Supplementary Fig. 2b), indicating that most PV interneurons expressed PSAM-GlyR. As illustrated in Fig. 3g-h, the pharmacogenetic inhibition of PV interneurons induced by local injection of the exogenous ligand PSEM-89S suppressed 92.6±9.7% (median=97%, n=5 mice) of spontaneous fIPSPs, from 6.58±3.5 to 0.45±0.66 events/s (n= 5 mice, p<0.05, two-sided Wilcoxon Mann-Whitney Test), suggesting that the recorded fIPSPs originated predominantly from PV interneurons.

**Figure 2:**
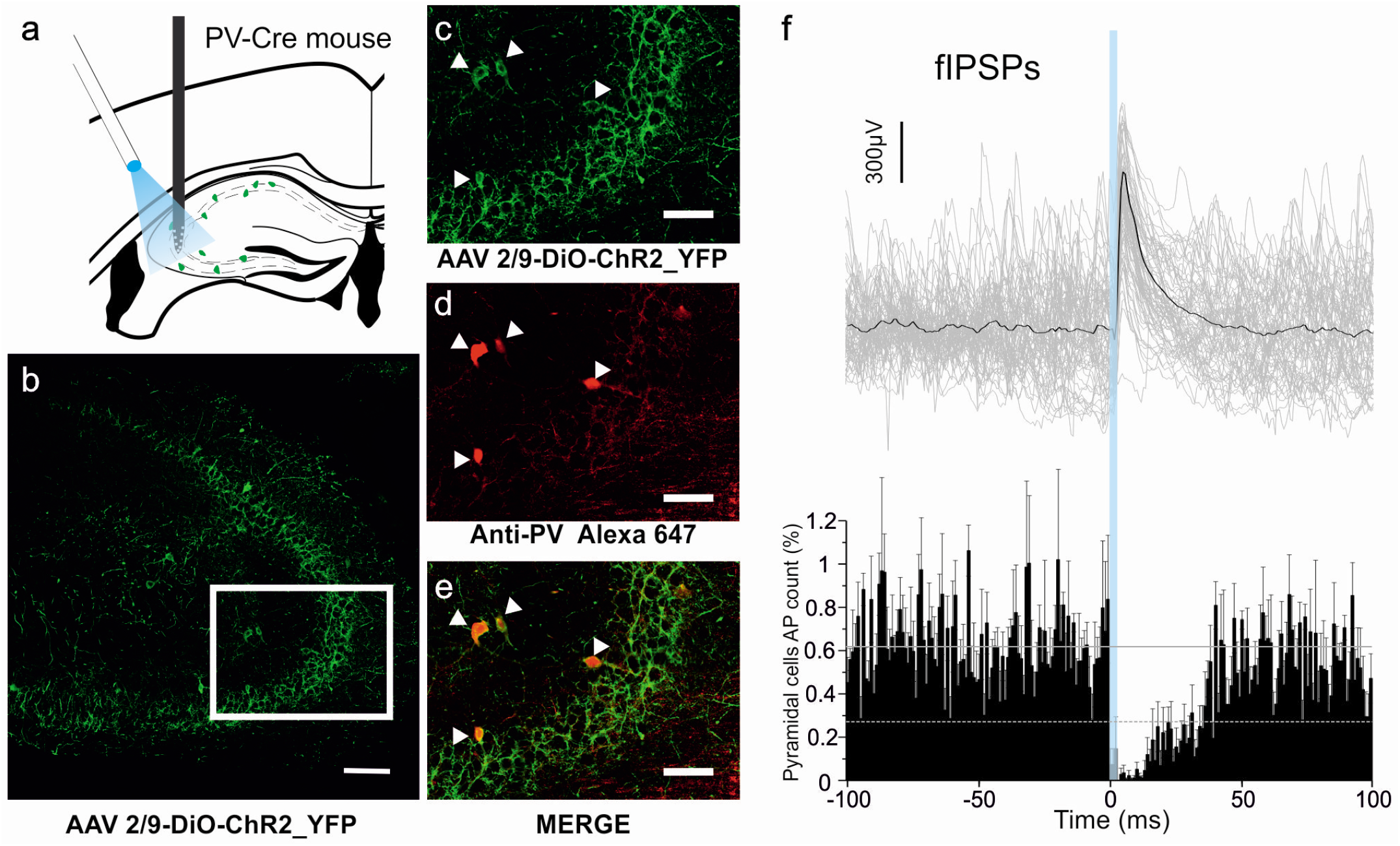
In vivo fIPSPs generated by optogenetic activation of PV interneurons. **(a)** Schematic representation of electrode and optic fiber positions in PV-Cre mice with AAV-driven ChR2 and E-YFP expression in CA3 PV interneurons. **(b-e)** Confocal stacks of E-YFP-positive (green) and PV-positive (red) immunostaining in the hippocampal CA3 region, **(b)** E-YFP immuno staining (**scale bar**, lOOμm). **(c-e)** higher magnification of the CA3a region delimited by the **white rectangle** in **b** (scale bar, 5Oμm). **(c)** E-YFP immuno-staining. **(d)** PV immunostaining, **(e)** Overlay. Note the co-localization on the 4 identified PV interneurons (**arrow-heads**). **(f)** Peri-event time histogram (mean ± SEM, n=5 mice, time bin 1 ms) between all the spikes discharged by putative pyramidal cells and optogenetically evoked (laser ON 2ms, blue bar) fIPSPs (reference), shown as superimposed (grey) and average (black) traces (n=6O from a single representative recording) on top, aligned and at the same time scale. Note the powerful time-locked inhibition of pyramidal cell firing in response to fIPSPs evoked by the optogenetic stimulation of PV interneurons.

**Figure 3:**
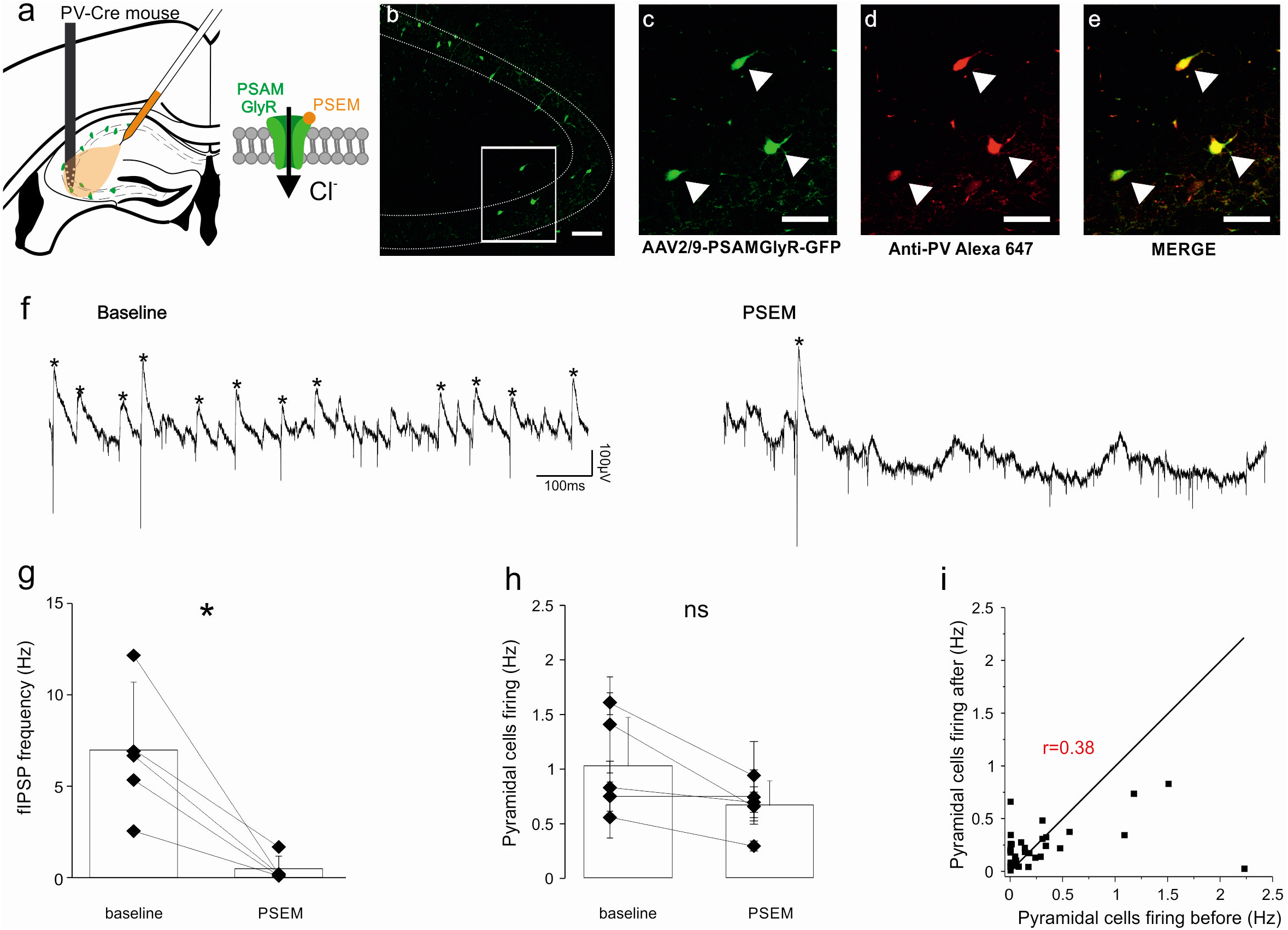
Inhibition of fIPSPs by pharmacogenetic inactivation of PV interneurons. **(a)** Schematic representation of electrode and injection site in PV-Cre mice with AAV-driven PS AM-GlyR and GFP expression in CA3 P V interneurons. **(b-e)** Confocal stacks of GFP-positive (green) and PV-positive (red) immunostaining in the hippocampal CA3 region, **(b)** GFP immuno staining (**scale bar**, 1 OOμm). **(c-e)** Higher magnification of the C A3a region delimited by the **white rectangle** in b (**scale bar,** 5Oμm). **(c)** GFP immuno-staining. **(d)** PV immunostaining, **(e)** Overlay. Note the co-localization of PV and PSAM-GlyR immuno-staining on the 3 labeled interneurons (**arrow-heads**). **(f)** Example trace of spontaneous wide-band (0. lHz-9KHz) field-recording activity in the CA3 pyramidal layer, in control (left, **baseline**) and in presence of the exogenous ligand PSEM-89S (right, **PSEM**). Note the drastic reduction in fIPSPs expression (*). **(g-h)** Summary plots (mean± SD, n=5 mice) of fIPSP frequency **(g)** and pyramidal population firing rate **(h)** in control (baseline) and in presence of PSEM (* p<0.05). **(i)** Individual pyramidal cells firing rates (n=42 putative pyramidal cells from 5 mice) in control (“before” corresponds to the lOmin period that immediately preceeds PSEM-89S injection) and in presence of PSEM (“after” is the period from 5 to l5min after PSEM-89S injection). Note that the pharmacogenetic blockade of PV interneurons with PSAM-GlyR redistributed individual firing rates without any significant change on global population firing activity.

Even though pharmacogenetic silencing of PV interneurons in the anesthetized mouse did not increase the global firing rate of the CA3 pyramidal cell population (Fig. 3i, 1.19±2.72Hz in control vs 0.96±1.04Hz in PSEM-89S, n=5 mice, NS, two-sided Wilcoxon Mann-Whitney Test; 1-way ANOVA, F(1)=1.51, NS), it did affect individual cells firing. As illustrated in Fig. S3, the correlation coefficient between individual cells firing rates in consecutive epochs of 10min indicated a remarkable stability, both in baseline (r=0.987, p<0.001) and after PV interneuron silencing (r=0.79, p<0.001). Application of PSEM-89S in transgenic PV-Cre mice not injected with the PSAM-GlyR viral construct did not affect global hippocampal activity (average global pyramidal firing rate, 0.6±0.21Hz in control vs 0.59±0.18Hz with PSEM-89S; fIPSP frequency of occurence, 6.72±1.71Hz in control vs 6.31±1.44Hz with PSEM-89S, n= 5 mice, Wilcoxon Mann Whitney ranks test, NS) or pyramidal cells individual firing rates (correlation coefficient between individual cells firing rates, r = 0.94, n=46 pyramidal cells from 3 mice). On the other hand, as illustrated in Fig. 3i, the firing rates before vs after PSEM-89S in PV-Cre mice expressing PSAM-Gly-R were hardly correlated (r=0.4, n=42 pyramidal cells from 3 mice), even after removing the values of the 4 outlier cells which average firing rates were above 1Hz in control condition (r = 0.35, n=38 pyramidal cells from 3 mice). Therefore, the global firing activity remained constant upon blockade of PV interneurons, while the average spiking rates of individual neurons were redistributed. This means that the decreased average spiking of some neurons was compensated by the increased average spiking of other neurons, so that the global population firing activity within the network remained unchanged. In other words, PV interneuron silencing induced a homeostatic redistribution of firing rates among pyramidal neurons within the CA3 circuit.

### Depolarizing and excitatory perisomatic GABAergic transmission in acute seizures in the drug-free mouse

The reversal potential of Cl-mediated GABAergic currents (E-GABA) is controlled by the passive movement and active transport of Cl^-^ through various ionic channels and transporters in the plasma membrane to reach the homeostatic set point for [Cl]i determined by local impermeant anions (44). During spontaneous activity, Cl^-^ ions flowing through GABA_A_ channels tend to shift the reversal potential of GABAergic currents toward membrane potential. Changes in external [K^+^] will also affect K-Cl cotransport and Cl^-^ gradient. E-GABA is therefore a very dynamic process that depends on ongoing fluctuations of membrane potential, GABAergic conductances and network activity. During seizures, GABA is massively released by interneurons while pyramidal cells are depolarized and therefore loaded with Cl^-^ until extrusion mechanisms restore an hyperpolarizing Cl^-^ gradient. It is expected that due to the time course of Cl^-^ extrusion, high Cl^-^ load and excitatory GABA might participate in a deficit of inhibition immediately after a seizure (16, 47). However, the time dynamics of spontaneous GABAergic synaptic transmission and Cl^-^ gradient through the interactions between the fluctuations of membrane potential and intrusion and extrusion systems, that have been studied in vitro, largely remain to be characterized in vivo. As illustrated in Fig. 4, taking advantage of fIPSPs as a non invasive index of the polarity of GABAergic currents in the CA3 pyramidal population, we have investigated the dynamics of perisomatic inhibition during acute interictal and ictal seizures in awake mice. Spontaneous fIPSPs in the head fixed condition had a higher frequency of occurrence than under urethane anesthesia (4.13±1.30 vs 8.57±2.05Hz, t(8)=3.63, p<0.01) but in both conditions they had similar rise times (1.81±0.23 vs 1.956±0.15 ms, t(8)=1.03, NS), decay time constants (3.65±0.45 vs 4.11±0.73ms, t(8)=1.04, NS) and amplitudes (206.4±9.35 vs 218.1±15.48μV, t(8)=-0.35, NS).

**Figure 4:**
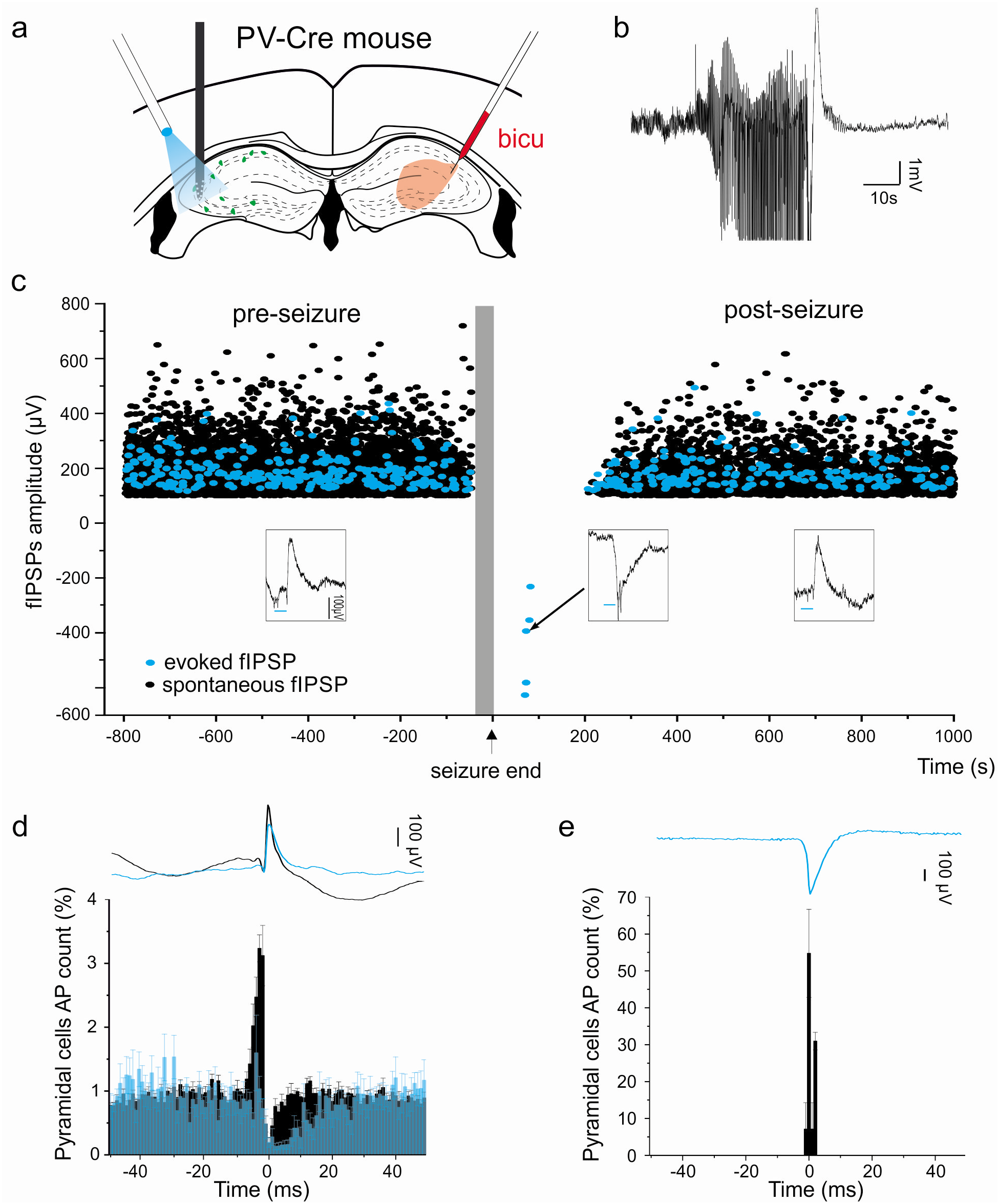
Transiently reversed fIPSPs and perisomatic GABAergic synaptic transmission following acute seizures. **(a)** Schematic representation of electrode and optic fiber positions in the left hippocampus of PV-Cre mice with A AV-driven ChR2 expression in CA3 P V interneurons, and contralateral injection of the convulsivant bicuculline. **(b)** Example trace (wide-band LFP) of a seizure as acute response to contralateral bicuculline injection. **(c)** Time course of individual fIPSPs amplitude (**black dots**, spontaneous, **blue dots**, evoked by optogenetic activation of PV interneurons, stimulation 2ms every 3s without interruption throughout the recording), relative to acute seizures induced by the contralateral injection of bicuculline (**vertical grey bar**, 3 seizures from the same animal, aligned by their ending time). **Insets**, representative individual field responses (optogenetically evoked fIPSPs; **horizontal blue bars**, laser ON) in control (**preseizure**), during the period right after the seizure (**reversed polarity**), or after recovery (**post-seizure**). Note transiently evoked spiking activity induced by fIPSPs of reversed polarity right after the seizure, and progressive recovery of fIPSPs amplitude. The absence of blue dots during and immediately after the seizure indicates the failures of optogenetically evoked response. **(d)** Peri-event time histogram (mean and SEM, time bin 1ms) between all the spikes discharged by putative pyramidal cells (n=7 mice) and spontaneous (in **black**, n=44435 events) or optogenetically evoked (in **blue**, n=7738 events) fIPSPs (reference) from the pre- and post-seizure periods, shown as average traces on top (spontaneous, **black**; evoked, **blue**) at the same time scale and aligned on peak. **(e)** Peri-event time histogram (mean and SEM, time bin 1ms) between firing activity (n=3 responding putative pyramidal cells from 2 mice out of 42 cells from 7 mice) and optogenetically evoked fIPSPs (reference) in the period right after the seizure (n=18 fIPSPs from 4 seizures with excited putative pyramidal cells, out of a total of 14 seizures), shown as average traces on top at the same time scale and aligned on peak. Note the clear time-locked excitation of pyramidal cell firing and reversed polarity of the fIPSPs evoked by the optogenetic stimulation of PV interneurons in the period right after the seizure.

In order to monitor both spontaneous and evoked fPISPs, PV-Cre mice were infected with the DIO-hChR2-EYFP construct in the CA3 region. Light stimulation with a locally positioned optical fiber provided evoked fIPSPs by direct stimulation of PV interneurons. Acute seizures were induced by a brief and focal intra-hippocampal injection of the convulsive agent bicuculline in the contralateral hemisphere. Electrophysiological recordings displayed both interictal activity and hippocampal ictal seizures, recognized as complex events lasting 30 to 40 seconds (37.4 ±9.3s, n=14 seizures from 7 mice), even though the animal did not show behavioural convulsions. Because the network can be durably altered by many repeated evoked seizures, even at times resulting in spontaneous seizures, we have decided to select only the first 2 or 3 evoked seizures of each animal, for homogeneity and to avoid the confound of a transition towards spontaneously recurring seizures (21). Optogenetic stimulations (2ms, 0.5Hz) were delivered without interruption throughout the recording session, but we observed that spontaneous and evoked fIPSPs were transiently interrupted upon ictal seizure onset. While in comparison to baseline, fIPSPs were not significantly affected between epileptic discharges, they transiently appeared with a reversed polarity after the termination of ictal seizures (latency to normal polarity, 119 to 406s, 224.5±259s, n=7 mice totalizing 14 seizures), suggesting transiently reversed polarity of GABAergic transmission after ictal seizures, but not after acute interictal discharges. Reversed fIPSPs often (n=9 over 14 seizures from 7 mice) entrained the discharge of postsynaptic target neurons, as illustrated by the short-latency action potentials evoked by fIPSPs in the traces of Fig. 4c and in the peri-event histogram between reversed fIPSPs and pyramidal cell firing activity in Fig. 4e (n=3 cells from a total of 42 putative pyramidal cells). The short latency (range 1-3ms) between fIPSP onset and pyramidal spike firing is compatible with direct excitation but not with previously described rebound from inhibition (38, 48, 49). No more fIPSP-related excitatory interactions were seen after the recovery of the polarity of fIPSPs. These results suggest that an ictal seizure can entrain transient depolarizing and excitatory actions of perisomatic GABAergic transmission in the CA3 circuit. Nevertheless, the expression of excitatory GABA, in terms of time-locked post-synaptic action potential firing, was restricted to less than 10% of the putative pyramidal cells identified, and confined to the postictal period devoid of spontaneous neuronal activity.

### Discrete excitatory perisomatic GABAergic transmission in a chronic model of epilepsy

Acute ictal seizures, as induced experimentally by the injection of the convulsive agent kainic acid (KA), can turn into chronic epilepsy after a silent period of up to several weeks. Previous studies on various models of epilepsy have suggested that reversed (excitatory) GABAergic transmission might be involved in the transition between acute seizures and chronic epilepsy (10–12, 14, 32, 50). This condition is of high clinical relevance because incident occurrence of isolated seizures in human is known to be a main factor for the development of chronic epilepsy after a latent period that can last from months to several years (51). Identifying the cellular mechanisms responsible for this transition would therefore potentially offer opportunities for preventive treatment against epilepsy.

We have evaluated the dynamics of perisomatic inhibition in the CA3 hippocampal region of anesthetized KA-treated mice during the latent period, one week after KA injection in the contralateral hippocampus, therefore at a stage when the animals are not yet epileptic (52). All KA-treated mice experienced acute status epilepticus. As illustrated in Fig. 5b and 6c, interictal discharges were present in all KA-treated mice (frequency of occurrence, 0.012±0.014Hz, n = 23 mice), but in none of the control animals (n = 18 mice). As illustrated in Fig. 5c, among all identified fIPSPs, none appeared of reversed polarity, whether spontaneous or evoked by the optogenetic activation of PV interneurons, in control (n=18 WT and 5 PV-ChR2 mice injected with Cre-dependant ChR2) or in KA-treated mice (n=8 WT and 15 PV-ChR2 mice injected with Cre-dependant ChR2), suggesting that GABAergic transmission remained globally hyperpolarizing at the network level. Nevertheless, because the polarity of fIPSPs represents the average response of all the local pyramidal targets of the discharging presynaptic interneuron(s), the possibility remained that a subpopulation of pyramidal cells might be depolarized or even excited by GABAergic transmission in spite of a globally hyperpolarizing fIPSP population response. We therefore evaluated the responses of individual neurons to spontaneous fIPSPs, and indeed identified a minority of neurons (n=4 out of 78 putative pyramidal cells from 3 KA-treated mice, among a total of 707 putative pyramidal cells from 23 KA-treated mice, n=0 out of 389 putative pyramidal cells from 18 control animals) that fired action potentials in response to GABAergic synaptic inputs. Individual examples of putative pyramidal cells displaying the three distinct types of observed responses to spontaneous fIPSPs (time-locked excitation, time-locked inhibition, or no clear change in net firing rate) are illustrated in Fig. 5. In fact, the most striking alteration of perisomatic inhibitory function in KA-treated mice is a loss of inhibitory control over most of pyramidal neurons. As illustrated in Fig. 6, the frequency of occurrence of fIPSPs was not significantly different in control and KA-treated mice (control, 7.99±3.39Hz, n=18 mice; KA-treated, 8.51±2.96Hz, n = 23 mice; t(39)=-0.52, NS), but only 45.1±30.6% of putative pyramidal cells were inhibited by spontaneous fIPSPs in KA-treated mice (median 38.8%, n=17 mice for a total of 572 cells) vs 71.7±12.9% in control animals (median 71.7%, n=18 mice for a total of 389 cells; MannWhitney test p<0.05). Putative pyramidal cells from KA-treated mice that were inhibited by fIPSPs presented the same strength and kinetics of inhibition as neurons from control mice (strength of inhibition: control, 90.6±1.9%, n=18 mice for a total of 280 inhibited pyramidal cells; KA-treated, 90.8±9.2%, n=23 mice for a total of 346 inhibited pyramidal cells, NS, MannWhitney test, p=0.87; time before recovery from inhibition: control, 7.92±1.85ms, n=18 mice for a total of 280 pyramidal cells; KA-treated, 8.22±1.73ms, n=23 mice for a total of 346 pyramidal cells; NS, MannWhitney test, p = 0.2). Overall, pyramidal population activity was about 50% lower in KA-treated mice than in control (control, 0.81±0.32Hz, n=18 mice for a total of 389 pyramidal cells; KA-treated, 0.39±0.23Hz, n = 23 mice for a total of 703 pyramidal cells; MannWhitney test, p<0.001), in line with the hypothesis that perisomatic inhibitory function is not the main determinant of quantitative population activity in the CA3 hippocampal circuit. In future studies, it might be interesting to investigate at various delays after an acute seizure episode whether the observed inter-individual heterogeneity in the fate of perisomatic inhibitory function across KA-treated mice is involved in the distinct phenotypic outcome of individual animals, differentially affected by the initial insult.

**Figure 5:**
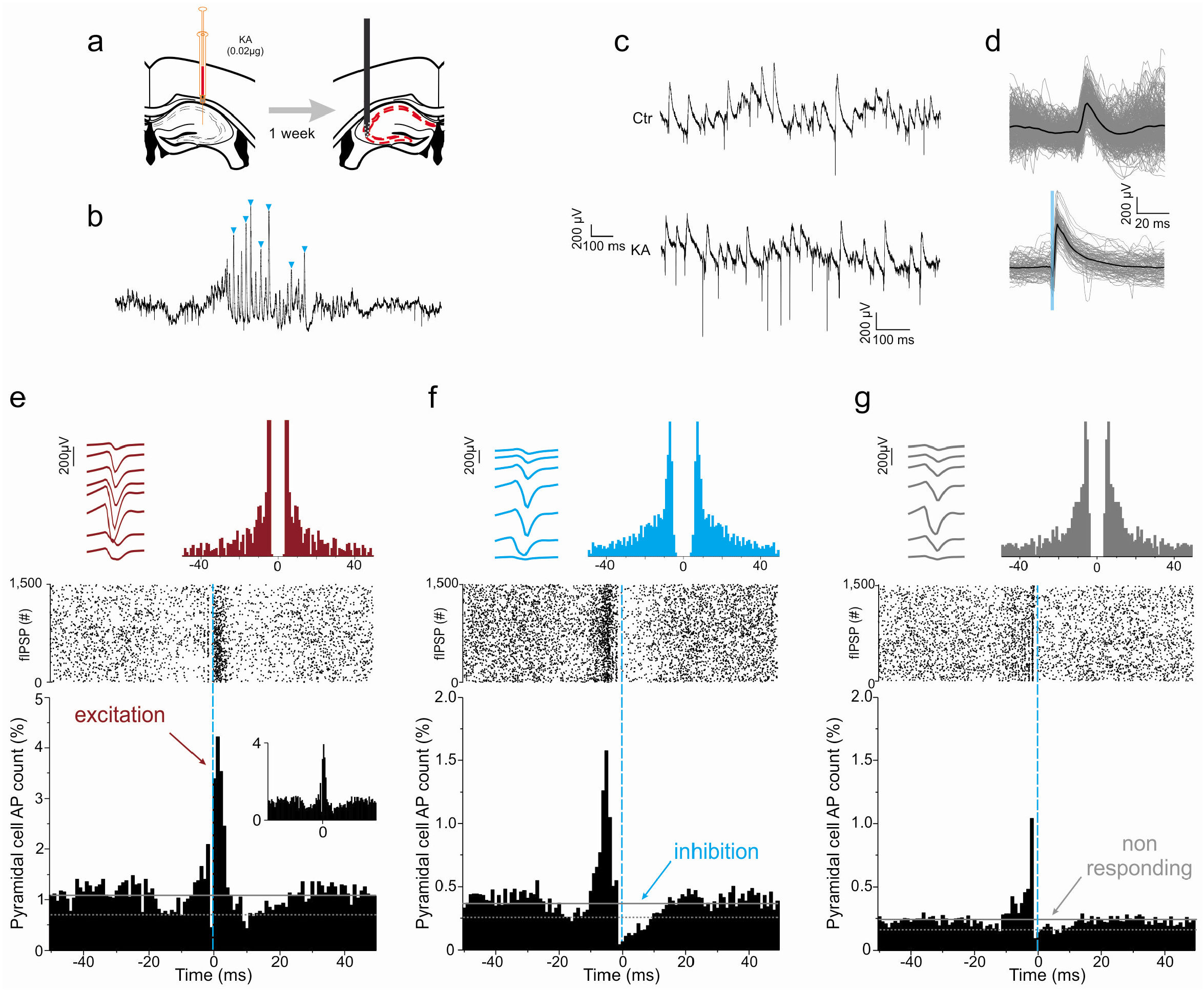
Normal fIPSP polarity but GABA-mediated time-locked excitation in head-fixed KA-treated mice. **(a)** CA3 neuronal activity was recorded from urethane-anesthetized WT mice one week after contralateral, intra-hippocampal KA injection. **(b)** Example trace (wide-band LFP) of interictal burst (interictal spikes, **blue arrow-heads**). **(c)** Example traces of spontaneous CA3 pyramidal layer activity in control (**top**) and KA (**below**) conditions, **(d)** Superimposed (**grey**) and averaged (**black**) traces of spontaneous (**top**) or optogentically evoked (**bottom**) fIPSPs in KA-treated mice. Note the presence of normal polarity fIPSPs in both conditions. **(e-g)** Individual examples of putative pyramidal cells displaying the three distinct types of observed responses to spontaneous fIPSPs in KA-treated mice (time locked excitation **(e)**, time-locked inhibition **(f)**, or no clear change in net firing rate (**g**, non responding)). **Upper left**, average waveform from the 8 channels of the recording electrode. **Upper right**, spike autocorrelogram. **Middle**, raster display of spike firing aligned on fIPSP ocurrence (vertical blue line, fIPSP peak). **Lower plot**, normalized peri-event time histogram (time bin, 1ms) of the neuron discharge (AP count) relative to fIPSP peak (reference, **vertical blue line**). **Inset** (e), similar peri-event time histogram but ignoring action potential bursts (ie all spikes with preceding inter spike interval < 10ms omitted from the histogram).

**Figure 6:**
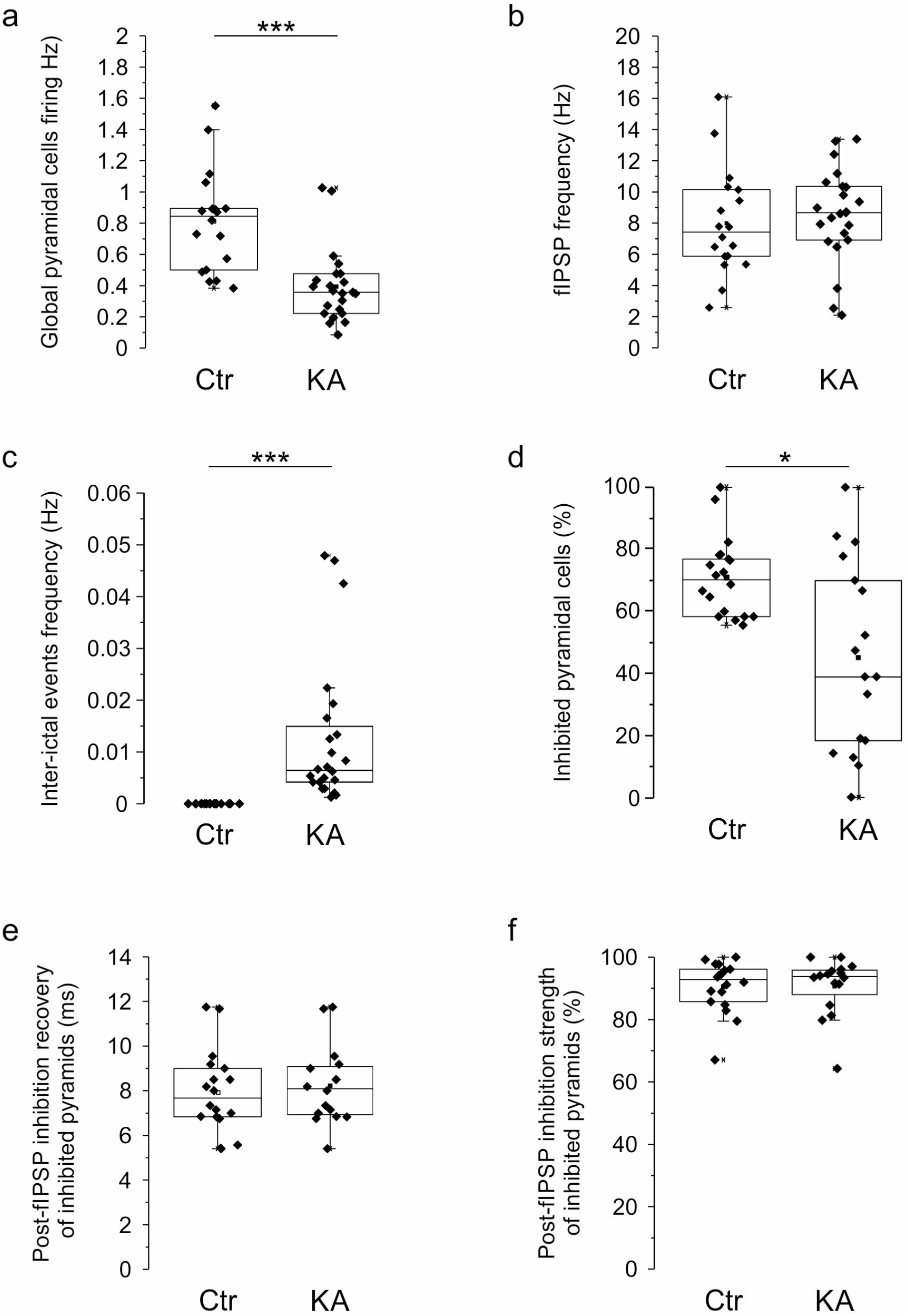
Loss of perisomatic inhibition in a majority of CA3 pyramidal cells during the latent, preepileptic period. Summary boxplots of various parameters of neuronal activity and perisomatic inhibition in CA3 putative pyramidal cells in control (Ctr, n=18 mice for a total of 389 cells) and KA-treated mice (one week post KA-injection,n=23 mice for a total of703 cells), *p<0.05, *** p<0.001 **(a)** average pyramidal firing rate (Hz), **(b)** frequency of fIPSPs occurrence (Hz), **(c)** inter-ictal event frequency of occurrence (Hz), **(d)** proportion of pyramidal cells inhibited by spontaneous fIPSPs (%), **(e)** latency to recovery (ms) from perisomatic inhibition for neurons that were actually inhibited by fIPSPs (last of post-fIPSPs time bin with firing rate below baseline - 2SD), and **(f)** strength of perisomatic inhibition (% of firing inhibition in the bin of minimum firing in the 4ms post-fIPSPs) for neurons that were actually inhibited by fIPSPs. Note the loss of perisomatic inhibitory control (ie non-responding cells) in the majority of pyramidal cells in KA-treated mice but the normal inhibition in responding neurons (inhibited by fIPSPs).

## Discussion

### CA3 fIPSPs to reveal the spontaneous dynamics of GABAergic transmission in vivo

The possibility that altered [Cl^-^]i regulation and functionally reversed GABAergic transmission (from inhibitiory to excitatory) might be involved in major pathologies such as epilepsy, autism spectrum disorders or schizophrenia, has recently raised considerable interest (2–5, 10–15, 19, 22, 23, 30, 31, 53, 54). However, a direct assessment of this hypothesis has been hindered by the technical difficulty of probing endogenous GABAergic synaptic function in vivo, because it implies to minimize perturbation of cytoplasmic content, membrane potential and natural synaptic GABAergic activity (18, 25, 55). The difficulties reside in both the recording methods and the activation of GABAergic transmission. Neuronal [Cl^-^]i is very labile and even intracellular recordings with sharp electrodes, which limit cytoplasmic wash-out, do not preserve membrane integrity, potentially altering resting potential and endogenous Cl^-^ gradient. Patch clamp (cell-attached) recordings are challenging in vivo, yield information about a single neuron at a time, and potentially affect neuronal resting potential (56), a major parameter for the function of GABAergic transmission. Imaging techniques (eg of calcium or chloride) are advantageous in terms of spatial resolution, but do not offer the time resolution of electrophysiological recordings and get invasive when applied to deep structures. Calcium imaging, in addition, does not detect depolarizations that do not reach the threshold of calcium channel activation, and hardly discriminates between depolarizing and excitatory effects that can both result in calcium increases, for example during a shunting depolarization not associated with action potential firing (57), or GABA_A_-induced calcium increases not mediated by changes in membrane potential (58). As for the activation of GABA_A_-R, neither the exogenous applications of GABAergic agonists, nor the electrical or optogenetic activation of GABAergic inputs used in previous studies (18, 27, 37, 38, 57, 59, 60), respect the dynamics of spontaneous synaptic transmission. For example, boosting Cl^-^ extrusion through KCC2 was recently shown to nicely prevent the prolongation of seizures induced by the optogenetic activation of PV interneurons, but had no effect on spontaneous, ongoing seizures (18). Therefore, in spite of a reversed Cl^-^ gradient and GABAergic polarity with important functional consequences if activated optogenetically, it remains unclear whether excitatory GABAergic transmission functionally participated in spontaneous ongoing seizure activity. Another caveat is that different types of interneurons interact with each other, such as the mutually inhibiting PV and SOM-expressing interneurons (39). Experimental manipulation of specific interneurons (eg PV) may therefore also influence seizure expression indirectly by the modulation of pyramidal cells’ GABAergic inputs from other interneurons. This illustrates the potential confound of interpreting the effects of GABAergic activation at the network level rather than getting a direct readout of monosynaptic, spontaneous GABAergic transmission.

On the other hand, previous work in vitro suggested that the direct postsynaptic response of hippocampal pyramidal cells to their spontaneously occurring perisomatic GABAergic inputs could be recorded extracellularly, as field-Inhibitory Postsynaptic Potentials (fIPSPs), and that the polarity of fIPSPs could be used as an index of Cl^-^ gradient and polarity of GABAergic transmission. Indeed, perisomatically projecting interneurons (basket and chandelier cells) have a densely arborised axon specifically confined to the pyramidal cell layer, and contact a large proportion of pyramidal cells within a restricted projection area (61, 62). Therefore, each action potential of the interneuron triggers the nearly synchronous release of GABA on hundreds of target pyramidal cells, which postsynaptic responses (IPSPs) sum up and can be readily identified from a local extracellular electrode (41–43), with a polarity reflecting that of the Cl^-^ gradient (41, 42). This approach is well suited to preserving endogenous [Cl^-^]i, can be performed in vivo, and opens the possibility to resolve the dynamics of Cl^-^ gradient and polarity of GABAergic transmission over the course of ongoing synaptic activity. A strong limitation however is that fIPSPs are the average response of multiple pyramidal cells and therefore lack the resolution of individual cells, so that changes in GABAergic polarity could be detected if they affect the whole neuronal population but may be missed if affecting only a subset of cells. Morevover, as mentioned before, GABAergic transmission involves both voltage and conductance changes, so that reversed Cl^-^ gradient and depolarizing GABA carry a complex combination of excitatory (through depolarization) and inhibitory (through increased conductance) influences (25, 40, 55, 57, 63, 64). As a consequence, determining the net postsynaptic effect of GABAergic transmission requires a direct readout of postsynaptic firing activity. For all these reasons, we now propose that the method of choice to probe Cl^-^ gradient and GABAergic synaptic function in vivo lies in combined multi-electrode recording and spike-sorting methods. While fIPSPs provide the timing and global (average) polarity of spontaneous perisomatic GABAergic synaptic events, the distributions (peri-event time histograms) of the spikes of individually identified neurons provide the net functional effect of GABAergic transmission, with single cell resolution.

### Qualitative vs quantitative control of CA3 pyramidal cell activity by perisomatic inhibition from PV interneurons

In line with previous in vitro evidence for the direct involvement of basket and chandelier cells in the generation of fIPSPs (41–43), we provide direct evidence that in vivo fIPSPs originate mostly from PV interneurons, because they were elicited by the optogenetic activation of PV interneuronal firing, and largely eliminated by the pharmacogenetic inhibition of PV interneuronal firing. Their pharmacological response to the GABA^A^ modulators zolpidem and diazepam also suggests that fIPSPs rather originate from perisomatically projecting basket cells than from axo-axonic or bistratified interneurons (45, 46). Perisomatic inhibition is considered to be functionally optimal for the control of neuronal discharge, with synapses strategically located on the soma and close to the site of action potential initiation (65, 66). PV basket and chandelier cells are thus considered to play a major role in the control of pyramidal cells firing through time-locked inhibition, presumably preventing excessive firing within the hippocampal circuit. However, in spite of the critical importance of inhibitory control over the recurrent CA3 circuit, the literature is strikingly missing direct reports of the dynamics of excitatory-inhibitory interactions in CA3, presumably because of the technical difficulties to identify perisomatic inhibitory events in vivo. Our experimental results confirm that perisomatic GABAergic transmission provides very powerful time-locked inhibition of most CA3 pyramidal neurons. An interesting observation is that the blockade of PV interneuron firing was responsible for a redistribution of pyramidal cell activity, but was not accompanied by any major quantitative change of activity at the network level. This suggests that in the CA3 hippocampal region, perisomatic inhibition may be involved in the control of the timing of pyramidal cells discharge rather than quantitative control of the network discharge. Previous reports are compatible with this hypothesis. In a previous study conducted in vitro, we have reported that fIPSPs were rather poorly recruited by network activity, suggesting that perisomatic inhibition should be more efficient in shaping the timing of spike flow than in limiting excitatory runaway recruitment and preventing the generation of hyper-synchronized discharges. Previous report that transgenic mice with specifically disrupted glutamatergic inputs to parvalbumin-positive interneurons displayed hippocampo-dependent spatial memory impairment but no epileptic phenotype are also in support of this interpretation (67, 68).

### Altered GABAergic perisomatic inhibition in epileptogenesis

One major advantage offered by the extracellular recording of fIPSPs is the possibility to evaluate the efficiency, and even the polarity, of perisomatic GABAergic transmission in a variety of conditions, including pathological conditions in which GABAergic transmission is suspected to be affected, or even reversed, due to altered neuronal Cl^-^ homeostasis. Cl^-^ gradient is dependent on a number of factors, including membrane potential, the distribution of intra and extracellular impermeant anions, osmotic pressure, ionic content of the extracellular milieu (eg [K^+^]), recent neuronal activity involving Cl^-^ currents, and the dynamics of pumps and transporters involved in maintaining physiological Cl^-^ gradient (44, 69). Our experiments have tested the possibility to detect the short and long-term consequences of acute seizures on Cl^-^ regulation and GABAergic transmission, which is particularly important to understand the mechanisms of epileptogenesis and identify therapeutic actions against the development of epilepsy. We report that transient hypersynchronized discharges induce a global reversal of neuronal Cl^-^ gradient and polarity of GABAergic transmission, shifting from inhibitory to excitatory for a duration of several tens of seconds before returning to normal polarity and efficiency. The strongly depolarized membrane potential induced by ictal seizures (depolarizing block) is not in favour of a depolarizing Cl^-^ gradient and excitatory GABAergic transmission. On the other hand, subsequent neuronal hyperpolarization is likely to occur due to the activation of Ca-dependent potassium channels activated by the massive Ca increase that takes place in neurons during the post-ictal depolarizing block. We hypothesize that the fIPSPs of reversed polarity that were evoked after ictal seizures occurred during such period of strong neuronal hyperpolarization. Accordingly, we did not observe any fIPSPs immediately upon termination of ictal seizures, and the transient reversal of Cl^-^ electrochemical gradient and excitatory GABAergic transmission were confined to the period devoid of spontaneous neuronal firing. In line with recent work demonstrating the prolongation of seizures by reversed GABAergic transmission upon the optogenetic activation of PV interneurons but not during spontaneous activity (18), our results highlight the importance of probing the polarity of spontaneous GABAergic synaptic transmission in addition to the responses evoked by experimental stimulations.

While this reversal of GABAergic polarity in acute seizures is readily seen as a reversed polarity of fIPSPs, we also report a more subtle effect during the course of epileptogenesis in the KA model of chronic epilepsy. One week after KA injection, the majority of pyramidal cells escaped inhibitory control by perisomatic GABAergic events. Besides, we observed that perisomatic GABAergic transmission provided time-locked excitation of pyramidal neurons in the hippocampus, but restricted to a minor subset of neurons, and we did not observe a reversed polarity of fIPSPs. Therefore, subtle alterations in the regulation of Cl^-^ homeostasis and GABAergic transmission already operate in the hippocampal circuit during the latent period that precedes the establishment of chronic epilepsy. Independently of seizure generation, deficient perisomatic inhibition might significantly affect neuronal processing and information coding. The functional consequences of defective GABAergic transmission, both in terms of coding and circuit dynamics, and in other periods, other models of epilepsy, and other pathologies, therefore remain to be investigated. From our recordings, we may hypothesize that excitatory GABAergic transmission may have a more limited contribution to CA3 hippocampal epileptogenesis than suspected from previous studies (10, 12, 15–17, 29, 31–33), that were not based on direct in vivo evidence. On the other hand, we can not exclude the decisive participation of excitatory GABA at different sites than where our recordings were performed. The exact contribution of excitatory GABA may depend on the exact brain area and identity of the affected neurons.

Finally, the extracellular detection of perisomatic IPSPs provides an invaluable tool to evaluate Cl^-^ homeostasis and inhibitory function, in physiological conditions and in situations in which alteration or even reversal of GABAergic transmission is hypothesized to occur. This approach opens new possibilities to evaluate the relevance of the excitatory GABA hypothesis in physiological development, autism spectrum disorders, schizophrenia, various forms of epilepsy and the time course of epileptogenesis, as well as to evaluate the potential of pharmacological agents to interfere with neuronal Cl^-^ gradient, as an effective therapeutic strategy against major brain diseases.

## Materials and Methods

### Animals

A total of 58 male adult mice (age 5-10 weeks, body weight 30-40g) were used in this study. Animals were kept on a 12h:12h light/dark cycle and provided with ad libitum access to food and water. Among 32 transgenic mice expressing the Cre-recombinase under the control of the Parvalbumin promoter (PV-Cre, Jackson Laboratory, B6;129P2-Pvalbtm1(cre)Arbr/J), 27 were injected with Cre-dependant Channelrhodopsin2 (ChR2) for optogenetic activation of PV interneurons with light pulses delivered from a laser source through an optic fiber, among which 15 received an intrahippocampal injection of the convulsive agent kaïnic acid (Sigma-Aldrich Co., St Luis, MO), and 5 with the Cre-dependant Pharmacologically Selective Actuator Module fused to the glycine receptor (PSAM-GlyR) for pharmacogenetic silencing of PV-interneurons with the exogenous specific ligand PSEM-89S(70, 71). Among 26 wild-type mice (B6129SF2), 8 received an intra-hippocampal injection of kainic acid, and the other 18 served as controls. All experimental procedures were performed in accordance with the EU directives regarding the protection of animals used for experimental and scientific purposes (86/609/EEC and 2010/63/EU), with the French law, and approved by the Ethical committee CEEA50 (saisines #10894, 10896 and 19354).

### Surgical procedures

The mice were anesthetized with isoflurane (1.5 - 5%), placed in a stereotaxic frame (David Kopf Instruments) and kept on a thermal blanket (Physitemp) to maintain their body temperature at 37.5°C. The eyes were covered with a protective liquid gel (Ocrygel). The scalp and periosteum over the dorsal surface of the skull were removed. **For virus injection** (optogenetic and pharmacogenetic constructions), a small craniotomy was executed at the coordinates of the left dorsal hippocampus (AP-1.6mm and L2.5mm relative to Bregma), and the injections (500nl measured using a Hamilton syringe, 2mm deep from the brain surface) performed (n=32 PV-Cre mice) using a thin glass capillary (40-80μm at the tip). After injection, the capillary was maintained *in situ* for about 10 min before being slowly removed. The scalp was then sutured and the animal housed individually for 3-6 weeks before electrophysiological recording. **For head-restrained experiments** (n=7 PV-Cre mice), reference and ground wires were implanted in the cerebellum and attached to a 2-pin connector, maintained together with a small custom-made horizontal stainless-steel bar anchored to the skull above the cerebellum with dental acrylic. A thin anchoring layer of dental acrylic (Super-Bond, Frapident) was applied on the exposed skull, except at specific locations in which a thin layer of bone was kept intact to allow for dorsal CA3 targeting with the recording silicon probe (left hemisphere, AP-1.6, L2.5, vertical insertion) and optical fiber (AP-1.6, L 3.2, insertion with an angle of 15° from vertical) as well as contralateral (right hemisphere) dorsal hippocampus targetting of the capillary for bicuculline injection (AP-1.6 L1.5, insertion 20° from vertical). **For the study of chronic epileptogenesis** (n=8 WT mice and 15 PV-Cre mice injected with Cre-dependant ChR2), a craniotomy was performed above the dorsal hippocampus (right hemisphere, AP-1.6, L1.5) for vertical insertion of a capillary and injection of kainic acid (0,2μg dissolved in saline, 500nl measured using a Hamilton syringe) at 1.8 mm from the brain surface. After injection, the capillary was maintained *in situ* for an additional 5 min before being slowly removed. The scalp was then sutured and the animal housed individually for 1 week before electrophysiological recording. As described in previous work (52, 72), 90% of the mice injected with KA experience an initial status epilepticus followed by a latent period (ie without seizures) of about 2 weeks before expressing chronic epilepsy. We can therefore consider that most of the animals we recorded (n=23 mice) were in a pre-epileptic state. Bicuculline, diazepam and kaïnic acid were purchased from Sigma-Aldrich Co., St Luis, MO.

### Optogenetic excitation and chemogenetic inhibition of PV interneurons

For **optogenetic** excitation of PV interneurons, PV-Cre mice were injected with an adeno-associated virus (AAV) bearing ChR-2 with a YFP reporter, pAAV-Ef1a-DIO hChR2 (E123T/T159C)-EYFP (gift from Karl Deisseroth, packaged into AAV serotype 9 from Addgene, plasmid # 35509, initial viral titer ≥1×10*13 gcp/ml, diluted four times in HBSS). Optogenetic activation (light wavelength 473nm, power max 10-14mW, 2ms pulses, 0.5Hz, driven by a train generator and current stimulator from Digitimer Ltd) was delivered through an optical fiber (diameter 200μm) connected to a diode-pumped solid-state (DPSS) laser (Shangai instrument, 100mW or IKECOOL Corporation). For **pharmacogenetic** inhibition of PV interneurons, we used an AAV bearing PSAM-GlyR with a GFP reporter, -rAAV-syn::FLEX-rev::PSAML141F,Y115F:GlyR-IRES-GFP (Addgene, gift from Scott Sternson, plasmid # 32481, packaged into AAV serotype 9, initial viral titer ≥10*13 gcp/ml, diluted four times in HBSS). PSAM-GlyR activation was performed by the injection of the specific ligand PSEM-89S (150-200nl, 15μM in saline, obtained from Scott Sternson, Janelia Farm Research Campus, VA, USA), pressure-ejected into the left hippocampus from a glass capillary (40-80μm at the tip).

### Head-restrained recordings

The mice (n=7 PV-Cre), previously implanted with head-posts, were maintained in the stereotaxic apparatus from their implanted fixation bars. Their body was inserted into a plastic tube (diameter approximately 5cm) to provide a reassuring confined environment and prevent excessive body movement. The animals readily habituated to head-fixed conditions. After initial training periods of a few minutes, restraint duration was gradually increased each day until the mouse would sit calmly for a period of roughly 1h. Electrophysiological recordings were performed after several days (at least five) of habituation. The dominant behavioral state in the head-restrained condition was awake immobility. On the day of recording, under brief isoflurane anesthesia (and body temperature control with a heating pad, 37.5°C), the craniotomies were completed and the silicon probe, optical fiber and injection-capillary inserted. After 1h recovery from anesthesia, and baseline recording, bicuculline was injected (100-200nl, 100μM in saline) into the right dorsal hippocampus to induce acute hippocampal seizures. At the end of the recording session, the animals were sacrificed and their brains removed for histological verification of electrode and optical fiber positions. The seizures induced in the head-restrained awake condition are most often not convulsive. In case of convulsion, the animal was immediately anesthetized with isoflurane and the experiment interrupted. This experimental aspect has been specifically submitted and approved by the local ethics comittee (EC50) and the final protocol validated by the French Government (saisine #19355).

### Multi-site extracellular recording in vivo

Extracellular recordings of spontaneous multi-unit activity and local field potentials were performed on mice either drug free in the head-restrained configuration (n = 7 PV-Cre mice) or anesthetized (n=25 PV-Cre mice and 26 WT mice) with urethane (1.7g/kg, IP) and a complement of ketamine/xylazine (respectively 6.6 and 0.66mg/kg, IM, repeated whenever necessary, usually every 30 to 60min). Animals were fixed in the stereotaxic apparatus (David Kopf instrument), the craniotomy was performed, the dura was gently removed, and a multi-site silicon probe (Neuronexus Technologies, either A1×32-Poly3-5mm-25s-177, 32 channels arranged on a single shank as 3 rows of vertically arranged staggered recording sites, vertical separation 25μm, or Buzsaki16, 2 shanks separated by 200 μm, 8 recording sites each separated by 10-20μm), covered with DiI (Molecular Probes) for post-hoc verification of electrode position, was inserted vertically through the neocortex until the pyramidal layer of the CA3a hippocampal region. The stratum pyramidale was recognized by the presence of multiple firing cells and fIPSPs, and confirmed with the post-hoc identification of DiI labeling. Recordings were performed using either Lynx-8 amplifiers (Neuralynx, gain x1000, bandpass filter 0.1Hz to 9KHz) and a digitizer (United Electronics, 14bits, A/D Gain x2, sampling rate 20KHz) controlled by a custom-made LabView (v7.1, National Instruments) program, or an integrated Digital-Lynx SX system (Neuralynx, Cheetah v5.0 software, 24bits, sampling rate 32KHz, bandpass 0.1Hz to 9KHz and range ±5mV for LFP analysis, as well as bandpass 1Hz to 9KHz and range ±1mV for spike sorting), and stored on a PC for offline analysis. The pyramidal layer was recognized electrophysiologically by the typical presence of fIPSPs and multi-unit firing. The recording started approximately 30 minutes after the insertion of the probe. Urethane was purchased from Sigma-Aldrich, ketamine (Imalgene 1000) from Merial and xylazine (Rompun 2%) from Bayer.

### Histological Processing

At the end of the recording session, the animal was transcardially perfused with 4% paraformaldehyde. Coronal sections (60μm-thick, cut using a VT1000S Leica vibratome) were blocked in 5% Bovine Serum Albumin (BSA) and 0,25% Triton X-100. After Tris-buffered saline washes, sections were incubated overnight at 4°C in 1% BSA, 0,25% Triton X-100 with mouse primary antibody against parvalbumin (1:200, Sigma-Aldrich Co., St Luis, MO) and rabbit antibody against GFP conjugated to Alexa Fluor 488 (1:2000, Thermo Fisher). After Tris-buffered saline washes, sections were then incubated with 1% BSA, 0,25% Triton X-100 and Alexa Fluor 647 conjugated goat antibody to mouse (1:500, Thermo-Fisher) 4h at room temperature. After Tris-buffered saline washes, sections incubated 10 minutes with DAPI (1:1000 Sigma-Aldrich Co., St Luis, and MO) at room temperature. After washing, sections were mounted and coverslipped on slides. Confocal stack images from the CA3a hippocampal region were acquired. The number of GFP and PV-immunopositive cell bodies was counted on the stacks with the IMARIS software (Oxford instruments). To quantify co-localization, individual GFP labeled interneurons were identified and subsequently scored for PV immunoreactivity. Scans from each sample were collected in 3 slices per animal (n=3 ChR2 injected and 3 PSAM-GlyR injected PV-Cre mice).

### Data analysis

Data were visualized and processed using NeuroScope and NDManager from the Neurosuite software(73) (http://neurosuite.sourceforge.net), and analyzed using Excel (Microsoft Office 2013), Origin (OriginLab, Northampton, MA) and MATLAB (MathWorks) built-in or custom-built procedures. The spikes were detected and extracted (SpikeDetekt), automatically clustered (KlustaKwik), and the resulting clusters manually verified and refined (KlustaViewa) using the KlustaSuite software(74) (www.cortexlab.net/tools). Beside multi-unit activity (MUA), only the clusters with a clear refractory period and typical bursting pattern of putative pyramidal cells were included in the analysis of neuronal firing (and referred to as pyramidal cells). The fIPSPs were detected with the MiniAnalysis software (6.0.3 Synaptosoft Inc.) from the raw signal using the following parameters (manually adjusted winthin the indicated ranges []): amplitude (μV), min=100, rise time (ms), min=1, max=[4 10], decay (ms), min=[1 2], max=[6 10], halfwidth (ms) min=[1 2], max=[6 10], and decay10-90-slope (μV/ms), min=25, max=[500 1500]. In results, the value taken for rise time is the time to peak (latency from onset to peak) while the decay time constant is the constant value of the exponential curve best fitting the decay of the event. Epileptiform events (inter-ictal spikes) were identified with Minianalysis as events of positive polarity and larger than 5 times the average amplitude of the detected fIPSPs. All detected events were verified by visual inspection. Peri-event time histograms display the time-distributions of neuronal activity relative to fIPSPs detected from the same electrode/shank. The number of spikes was counted within time-bins of 1ms around reference events (either fIPSP-peak or optogenetic light-stimulation onset), and normalized by the total number of spikes within the histogram. Baseline activity was computed as the mean (and SD) of values between −50 and −30ms. Pyramidal cells were considered as inhibited by fIPSPs if their firing rate decreased and remained below that of [baseline - 2SD] from 0 to at least 4ms after fIPSP-peak. The strength of inhibition was then quantified as the percentage of firing inhibition (relative to baseline) in the inhibitory trough (bin of minimum firing in the 4ms post-fIPSPs), and the duration of recovery as the latency to the last of post-fIPSPs time-bin below [baseline - 2SD]. Only cells with [baseline - 2SD] > 0 were considered to have high and stable enough firing for the possible detection of potential fIPSP-mediated inhibition. Pyramidal cells were considered as excited by fIPSPs if their firing rate increased above [baseline + 3SD] within 4ms after fIPSP-peak. Because bursts of action potentials (short interspike intervals) which first spike would evoke a fIPSP would potentially produce a spurious post-fIPSP peak in the peri-event histogram that may be wrongly interpreted as pyramidal cell excitation, we also computed the peri-event histograms of putatively excited pyramidal cells excluding all the spikes preceded by an action potential by less than 10ms. The post-fIPSP peak-firing was still present in all the tested putatively excited pyramidal cells, excluding the contribution of misinterpretation due to burst firing. Pyramidal cells were considered as non-responding if they were neither inhibited nor excited by fIPSPs. In the quantification of the proportion of non-responding pyramidal cells in KA-treated mice, we included only the animals with at least 8 pyramidal neurons with enough spikes for peri-event time histograms (ie [baseline - 2SD] > 0). All statistical analyses were performed in Matlab (MathWorks) or Origin (OriginLab). All tests were two-tailed unless otherwise indicated. Our data did not follow normal distributions, therefore the non parametric Wilcoxon Mann-Whitney test was used. Results are displayed as mean ± s.e.m. unless indicated otherwise.

## Acknowledgements

We thank Roustem Khazipov, Rosa Cossart, Valérie Crepel, Gilles Huberfeld, Tarek Deeb and Isabel Del Pino for useful comments and discussions on a previous version of the manuscript, Delphine Gonzales, Nathalie Aubailly, and all the personnel of the Animal Facility of the NeuroCentre Magendie for animal care, Katy Lecorf for her help regarding virus preparation, the UMR5293 vector core facility for producing AAV vectors, and the laboratory of S.M. Sternson (Howard Hughes Medical Institute) for providing the chemical compound PSEM-89S.

## Author contributions

XL and HH performed pilot experiments and initiated the study. XL conceived and designed the experiments. OD and XL planned the study. OD, AFGDS, AB and XL performed the experiments. OD and XL analyzed the data. AF contributed with lab and breeding space, and a fraction of the running expenses. OD, AB and XL wrote the paper.

## Data availability

The data shown in the paper will be made available upon reasonable request to the corresponding author.

## Code availability

The custom code used for analysis will be made available upon reasonable request to the corresponding author.

---

**Figure S1:**
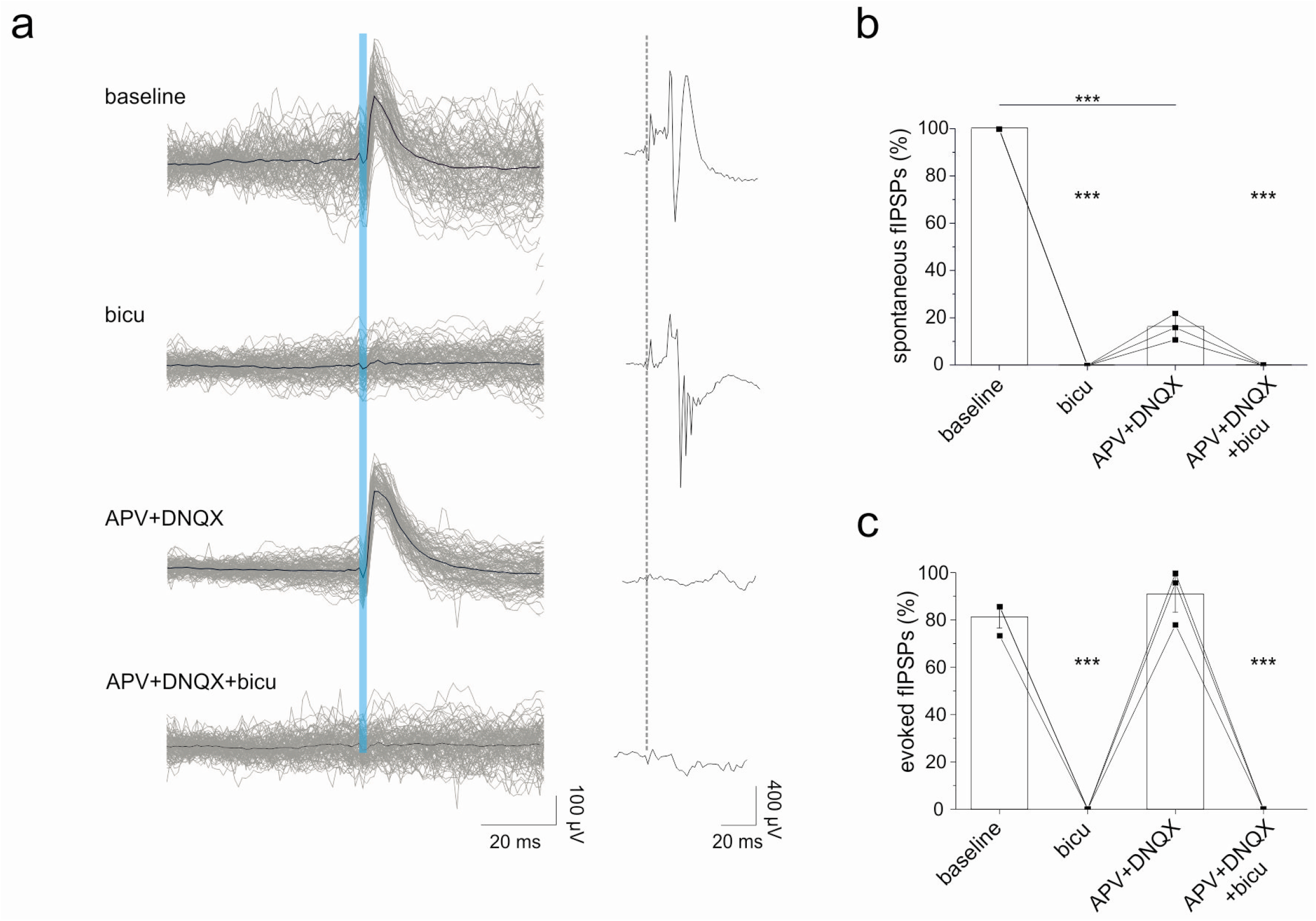
Inhibition of fIPSPs by pharmacological blockade of GABA_A_ but not of glutamatergic receptors. **(a)** Example traces of fIPSPs (**left, grey**, superimposed, **black**, average) evoked by the optogenetic stimulation of PV-intemeurons (**vertical blue line**, laser ON), and of the LFP response evoked on the same recording electrode by the electrical stimulation of the contralateral CA3 region (**right**). Note that the GABA_A_ antagonist bicuculine (**bicu**) blocked evoked fIPSPs and increased the electrically induced population response to multiple spikes, while the glutamatergic antagonists APV and DNQX abolished the electrically induced population response but not the evoked fIPSPs. **(b-c)** Sumary plots (mean ± SD, n=3 mice) of the effects of bicuculine and APV and DNQX on the frequency of occurrence of spontaneous fIPSPs (**b**, normalized to baseline frequency) and on the success rate of evoking fIPSPs by the optogenetic stimulation of PV interneurons (c). *** p<0.001

**Figure S2:**
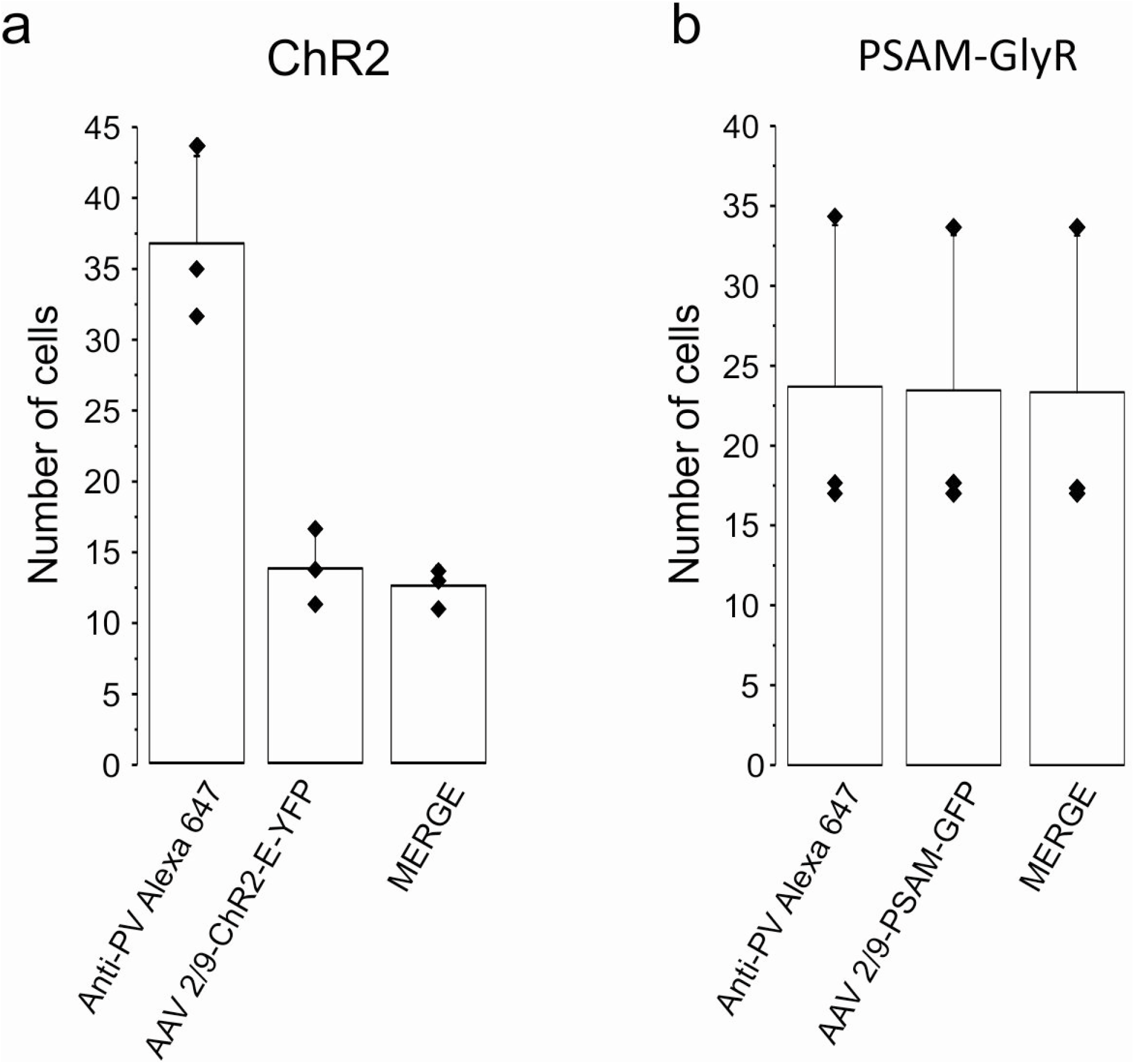
Immuno-histological control of ChR2 and PSAM-GlyR expression in PV interneurons. **(a)** Summary plot (mean ± SD, n=3 mice) of the number of cells immunopositive for PV (anti-PV Alexa 647), for ChR2 E-YFP, or for both (merge). Note that only a fraction of PV cells express ChR2 but that virtually all cells expressing ChR2 are PV cells, indicative of high specificity of ChR2 expression for PV cells. **(b)** Summary plot (mean ± SD, n=3 mice) of the number of cells immuno-positive for PV (anti-PV Alexa 647), for PSAM-GlyR-GFP, or for both (merge). Note that almost all PV cells also express PSAM-GlyR and that virtually all cells expressing PSAM-GlyR are PV cells, indicating high specificity and high coverage of infection for PV cells.

**Figure S3:**
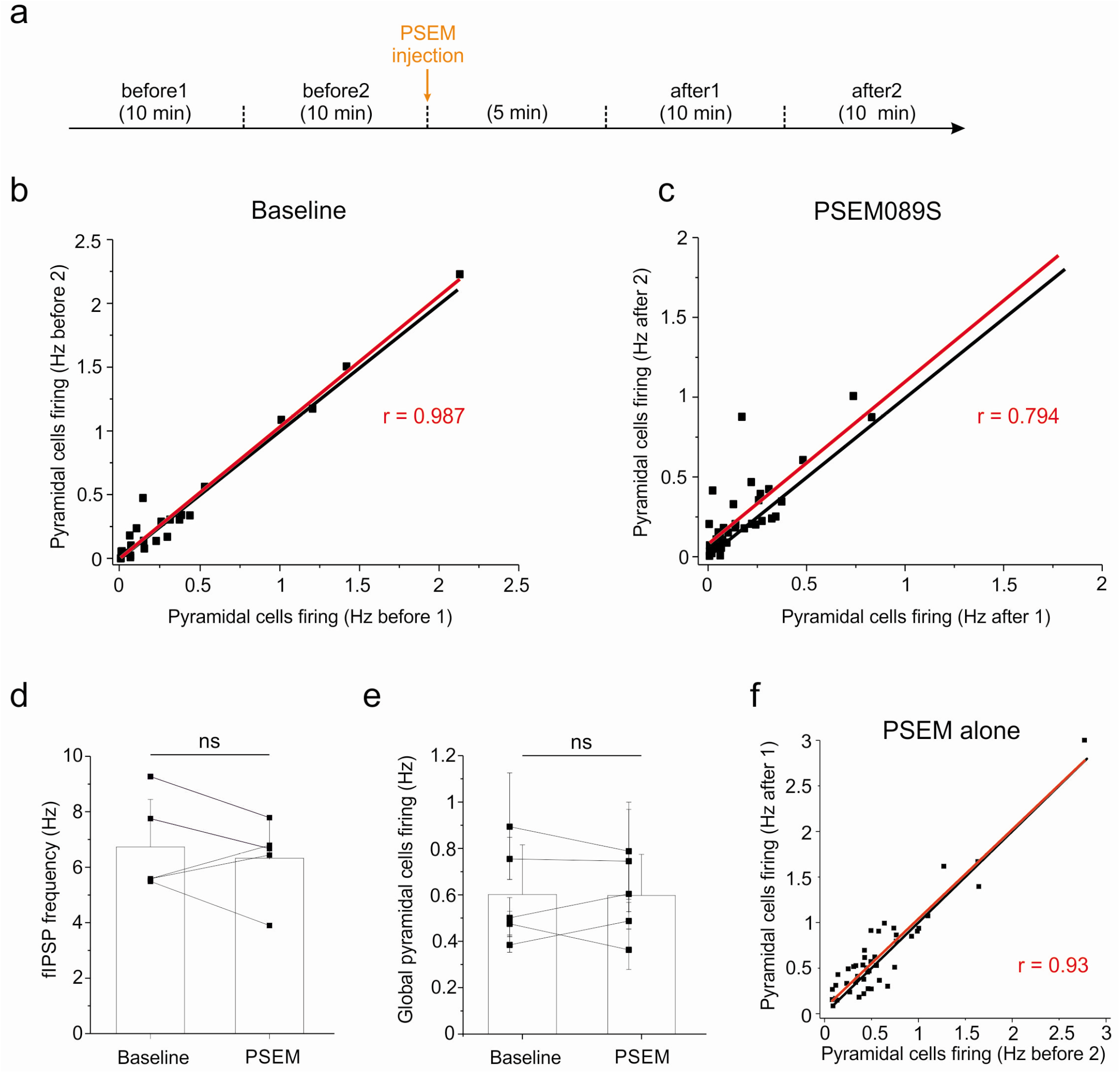
Redistribution of pyramidal cells firing rates after the pharmacogenetic blockade of PV interneurons. Comparison of discharge frequency of individual pyramidal cells (n = 42 neurons from 5 mice) in two successive recordings of lOmin, either both before (**b, baseline**) or both after (**c, PSEM-89S**) the injection of PSEM-89S, as depicted in the **timeline (a)**. Note high correlation values (**r**) in both baseline and PSEM-89S, indicating stable activity among CA3 pyramidal cells. **(d-e)** Summary plots (mean ± SD, n=5 mice) of flPSP frequency **(d)** and pyramidal population firing rate **(e)** in control (baseline) and in presence of PSEM-89S in WT mice (ie not expressing PSAM-GlyR). (1) Individual pyramidal cells firing rates (n=46 putative pyramidal cells from 5 mice) in control (“before2” corresponds to the lOmin period that immediately preceeds PSEM-89S injection) and in presence of PSEM (“afterl” is the period from 5 to l5min after PSEM-89S injection) in WT mice (ie not expressing PSAM-GlyR).

